# Non-invasive optoacoustic imaging visualizes exercise-induced dermal revascularization in obese mice

**DOI:** 10.1101/2024.03.26.586767

**Authors:** Shan Huang, Hailong He, Robby Z. Tom, Sarah Glasl, Pia Anzenhofer, Andre C. Stiel, Susanna M. Hofmann, Vasilis Ntziachristos

**Author notes:** Corresponding authors: Susanna M Hofmann Vasilis Ntziachristos. Equal contribution (Shan Huang, Hailong He).

## Abstract

Microcirculatory dysfunction in dermal (dWAT) and subcutaneous white adipose tissue (scWAT) of obese humans may predict cardio-metabolic disease progression. *In-vivo* visualization and monitoring of microvascular remodeling in these tissues remains challenging. We compared performance of multi-spectral optoacoustic tomography (MSOT) and raster-scanning optoacoustic mesoscopy (RSOM) in visualizing lipid and hemoglobin contrast in scWAT and dWAT of diet-induced obese (DIO) mice undergoing voluntary wheel running. MSOT quantitatively visualized lipid and hemoglobin contrast in fat depots at early stages of DIO. RSOM precisely visualizes microvasculature with quantitative readouts of skin layer thickness and vascular density in dWAT and dermis. Combination of MSOT and RSOM resolved exercise-induced morphological changes in microvasculature density, tissue oxygen saturation, lipid and blood volume content in dWAT and scWAT. Combination of MSOT and RSOM precisely monitor microcirculatory dysfunction and intervention response in dWAT and scWAT of DIO mice. Our findings lay out the foundation for future clinical studies using optoacoustic-derived vascular readouts from adipose tissues as a biomarker for monitoring microcirculatory function in cardio-metabolic disease.

## Introduction

Vascularization regulates adipose tissue function (Cao, 2010, Rupnick, Panigrahy et al., 2002, Ye, 2011). Vascular function in adipose tissue is a key factor for the development of metabolic disease and thus blood vessels are a promising target for overcoming metabolic perturbations associated with obesity (Cao, 2010, Graupera & Claret, 2018, Martinez-Santibanez, Cho et al., 2014, Sun, Kusminski et al., 2011, Trayhurn, 2013). As a result, the study of the interplay between vascularization and adipose tissue has attracted increasing attention (Hutchings, Janowicz et al., 2020, Laschke, Spater et al., 2021). Non-invasive examination of vascular function *in vivo* in adipose tissues may be suitable as a biomarker for disease monitoring or for examining treatment efficacy in obesity and related metabolic diseases.

Immunostaining of isolated tissue samples is currently used for *ex vivo* studies of vascularization in adipose tissues in both humans and mice (Pasarica, Sereda et al., 2009, Seo, Riopel et al., 2019, Tong, Zhang et al., 2020, Xue, Lim et al., 2010). However, this approach is not suitable for longitudinal observations as it requires multiple tissue biopsies or sacrifice. Thus, a non-invasive imaging tool that can visualize and quantify lipids and blood constituents simultaneously or allow repeated assessments of vascular function in a pathological or therapeutic context would be essential for longitudinal monitoring of disease progression and treatment response. Even though non-invasive imaging methods such as Magnetic Resonance Imaging (MRI) (Gronemeyer, Steen et al., 2000, Hu & Kan, 2013), Computed Tomography (CT) (Mrzilkova, Michenka et al., 2020, Schlett & Hoffmann, 2011), and ultrasonograpy (Bazzocchi, Filonzi et al., 2016, Clerte, Baron et al., 2013) can visualize the change in adipose tissue volume under physiological, pathological and therapeutic conditions, they require contrast agents for measuring microvasculature or blood volume parameters in adipose tissue, challenging *in vivo* and disseminated applications.

The multi-faceted functions of dermal white adipose tissue (dWAT) have recently drawn much research attention and may also serve as prediction markers for metabolic disease progression (Alexander, Kasza et al., 2015, Driskell, Jahoda et al., 2014, Guerrero-Juarez & Plikus, 2018, Zhang, Shao et al., 2019). It has been observed that vessel density in adipose tissues decreases under obese conditions in mice and humans(Graupera & Claret, 2018, Trayhurn, 2013). This phenotype can be rescued by exercise in mice (Driskell et al., 2014, Kolahdouzi, Talebi-Garakani et al., 2019, Min, Learnard et al., 2019, Stanford, Middelbeek et al., 2015a, Stanford, Middelbeek et al., 2015b). Studies that used rodent obesity models to demonstrate the vasculature dysfunction in adipose tissues in obesity(Shimizu, Aprahamian et al., 2014, Stanford et al., 2015b, Voros, Maquoi et al., 2005), or its potential as a therapeutic target(Brakenhielm, Cao et al., 2004, Cao, 2010, Sung, Doh et al., 2013, Xue, Xu et al., 2016), employed end-point assays such as staining or *in vitro* assays to analyze vessel functions. However, such invasive tissue interrogation is not suitable for monitoring the vascular changes *in vivo* and longitudinally. Therefore, the effects of obesity-induced vascular dysfunction and the potential for exercise to rescue this phenotype have not been demonstrated in live animals.

In-vivo and longitudinal observations could be enabled by optoacoustic imaging. Optoacoustic visualization can interrogate tissues at the microscopic (∼1mm depth), mesoscopic (<1cm depth) or macroscopic level (>1 cm depth) and simultaneous deliver anatomical, functional and molecular contrast (Omar, Aguirre et al., 2019, Taruttis, van Dam et al., 2015). Within the family of optoacoustic imaging implementations, Multispectral Optoacoustic Mesoscopy (MSOT) operates at the macroscopic regime and can separate spectral contributions of tissue chromophores such as lipids or oxygenated and deoxygenated hemoglobin by multi-wavelength illumination (Ntziachristos & Razansky, 2010). MSOT has been further employed to visualize brown adipose tissue (BAT) and white adipose tissue (WAT) *in vivo* and to record BAT activation under cold exposure or drugs in animals and humans (Li, Schnabl et al., 2018, Reber, Willershauser et al., 2018) or the distribution of lipoma and its vascularization (Buehler, Diot et al., 2017). At the mesoscopic regime, Raster-scanning optoacoustic mesoscopy (RSOM) reaches depths of several mm in tissue with resolutions in the 10-30 µm resolution (He, Schönmann et al., 2022, Omar et al., 2019), much higher than that of MSOT, which is in the range of no less than 100 µm (Ma, Taruttis et al., 2009). RSOM has been used to study microvasculature in epidermis and dermis in human skin or assess skin microvasculature changes in patients with diabetes (He, Fasoula et al., 2023).

Herein, we explore for the first time to our knowledge the suitability of optoacoustic methods in the study of adipose tissue and its vascularization and investigate the relative performance of MSOT and RSOM in dWAT imaging. We demonstrate that MSOT can non-invasively image and quantify lipids and blood content in adipose tissues in obese and non-obese mice and as a function of exercise. We recapitulate findings of vascular dysfunction in interscapular BAT (iBAT) and scWAT in diet-induced obesity (DIO) *in vivo* and observe a previously undisclosed decrease in vessel density associated with dWAT in obesity, which can be rescued by exercise. The MSOT findings are confirmed by histology and RSOM and suggest a new role of dWAT vessel density as a possible biomarker for metabolic disease monitoring. In contrast to scWAT that is located deeper in tissue, dWAT sits below the dermis and can be reached by RSOM at a higher resolution than MSOT. Overall, our findings suggest optoacoustic imaging as a suitable method for assessing tissue lipids, vasculature and blood content without contrast agents or endogenous labels. Therefore, the optoacoustic method may facilitate longitudinal and *invivo* animal research and drug development and has the potential to be clinically applied for *in vivo* monitoring of metabolic biomarkers.

## Results

### MSOT of lipid, total blood volume and tissue oxygenation in scWAT and iBAT of lean male mice

MSOT (Figure 1A) identified three distinct fat depots, iBAT, scWAT and dWAT in the neck region of male mice (Figure 1A, lower panel). To verify that MSOT can distinguish BAT from WAT, we acquired spectral data in the 700nm to 960nm range from regions of expected iBAT and scWAT (Figure 1B) and then unmixed the signals determining the lipid, oxygenated hemoglobin and deoxygenated hemoglobin content. Already from a qualitative inspection of the mean spectra (mean over area and N=5 mice) between 700 nm and 880 nm, the lipid peak at 930 nm is relatively more prominent in the case of scWAT suggesting higher lipid content. In contrast relative signals of the spectral region corresponding to hemoglobin absorption was more pronounced for iBAT suggesting higher vascularization relative to lipid. Using the spectral data, we unmixed the lipid and blood contents in scWAT and iBAT. As suggested by the spectra shown in Figure 1C, scWAT contained significantly more lipid than iBAT, while iBAT exhibited a higher total blood volume (TBV) than scWAT (Figure 1D). Tissue oxygenation (sO_2_) readout measured by MSOT revealed that sO_2_ rates between iBAT and scWAT were similar (Figure 1E).

**Figure 1.**
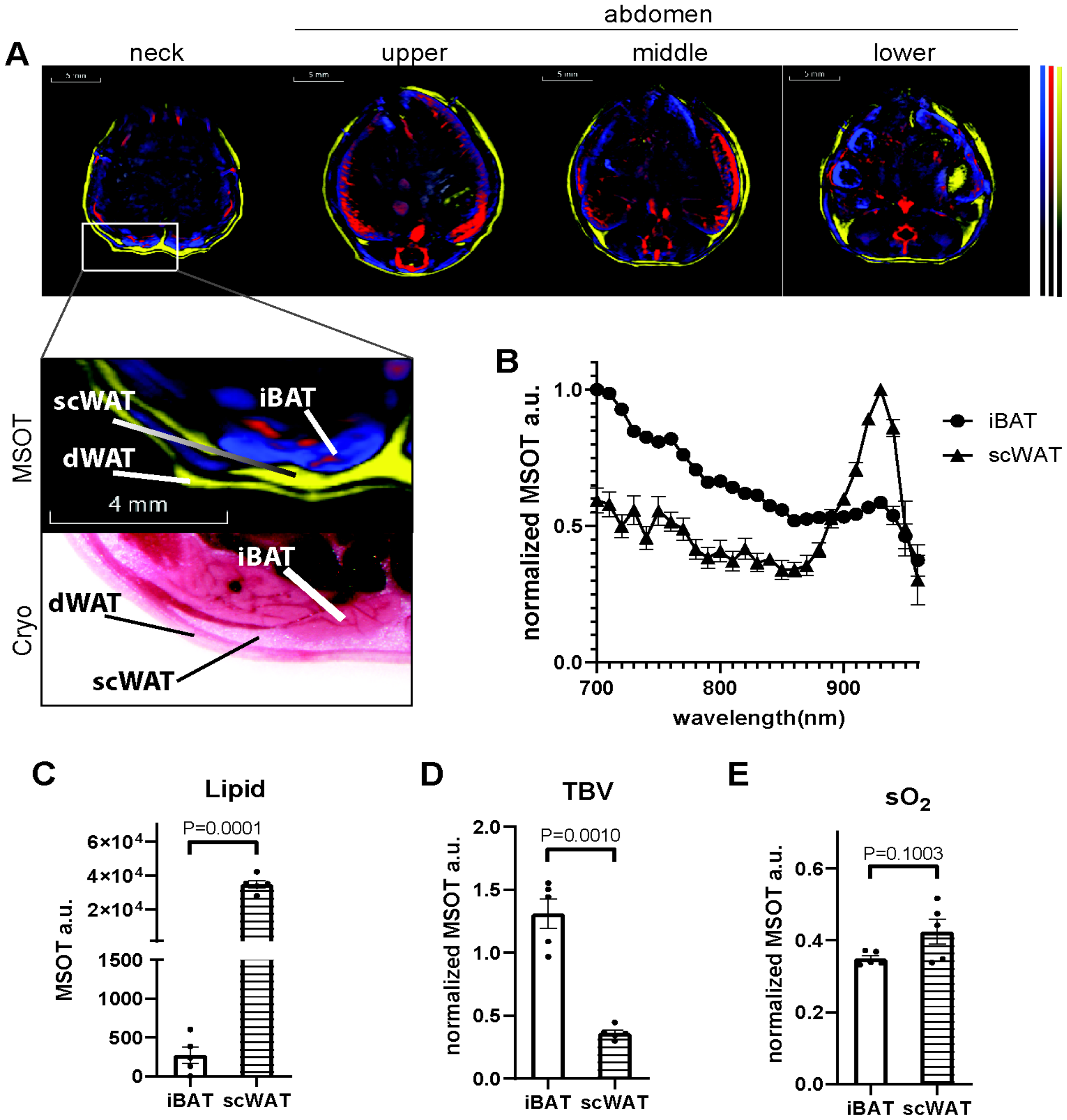
MSOT imaging of brown adipose tissue (BAT) and white adipose tissue (WAT) in vivo. A. Reconstructed MSOT image (800nm) with linear unmixing data in neck, upper, middle and lower abdominal area. Higher magnitude images are showing neck region with anatomical reference from cryo section image. Unmixing result: blue for Hb (deoxy-haemoglobin), red for HbO2 (oxy-haemoglobin), yellow for lipid. The color bar shows the color coding of MSOT a.u. from minimum to maximum (bottom to top). B. For visualization purposes spectra are normalized to maxima. Spectra of interscapular brown adipose tissue (iBAT) and subcutaneous white adipose tissue (scWAT). n = 5. C. Unmixing result of lipid from iBAT and scWAT, n = 5. D. Total blood volume (TBV) results from iBAT and scWAT, n = 5. E. Tissue oxygenation (sO2) results from iBAT and scWAT, n = 5. dWAT: dermal white adipose tissue.

### MSOT of lipid and blood content in murine fat depots at the onset of DIO

Following the MSOT assessment of adipose tissue in male mice fed with chow, we studied whether MSOT could monitor obesity-mediated pathological changes in adipose tissues in a DIO mouse model. Male mice were fed high fat diet (HFD) or chow for up to 6 months. The body weight of HFD-fed mice was significantly higher than chow-fed mice (Figure S1A). The weight of gonadal fat (gWAT), a visceral fat depot, inguinal fat (ingWAT) and a scWAT depot were all increased in HFD fed-mice, confirming a DIO phenotype (Figure S1 B-D). Consistently, MSOT measurements revealed an increase in lipid content in both iBAT and scWAT upon HFD feeding compared to chow feeding (Figure 2A). To enhance the visualization and relative comparison of spectral features, we normalized all tissue spectra to the highest optoacoustic signal acquired in the 700–960 nm range (Fig. 2B and 2C). Qualitative inspection of the spectra, especially the relative contributions of regions from lipid absorption (930 nm) and hemoglobin (680-900 nm), of iBAT from HFD-fed mice showed a much more prominent lipid peak at 930 nm compared to chow-fed mice, indicating a higher lipid content (Figure 2B). In contrast, the change in scWAT spectra caused by DIO was less obvious (Figure 2C). After spectral unmixing, the lipid content readout from iBAT showed significant increase in DIO mice compared to lean mice, indicating an ectopic accumulation of lipid in the tissue (Figure 2D, E). This observation was consistent with our histology findings showing larger fat vacuoles in iBAT from DIO mice, as compared to lean mice (Figure S2A). By quantifying the coverage of fat area from histological images, we found that iBAT from DIO mice has higher fat coverage than that from lean mice (Figure S2B). Compared to iBAT, we observed a less obvious change in the spectral composition in scWAT upon HFD feeding (Figure 2F). The total blood volume in both iBAT and scWAT were all decreased in DIO mice (Figure 2G, H, I). These findings were consistent with findings from other studies using end point *ex vivo* methods (Graupera & Claret, 2018, Trayhurn, 2013). Even after one month of HFD feeding, similar results were obtained using MSOT in the same cohort, which indicates that vessel function was already altered, albeit to a lesser extent (Figure S3).

**Figure 2.**
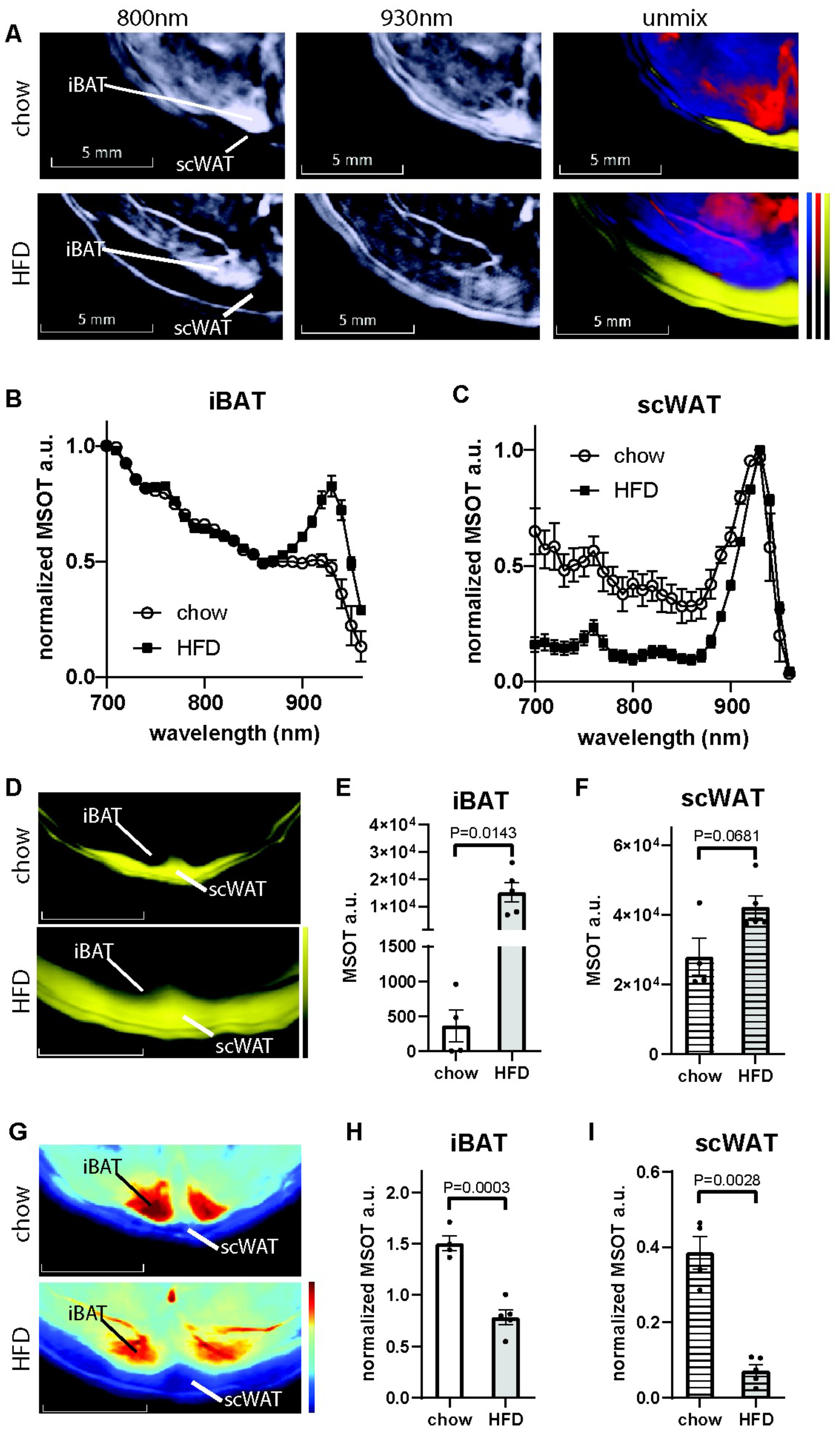
Comparison of blood and lipid content of iBAT and scWAT between healthy and DIO male mice using MSOT imaging. A. Reconstructed MSOT image (800 nm) with linear unmixing data of Hb, HbO_2_, and lipid from chow and high fat diet (HFD)-fed mice. The color bar shows the color coding of MSOT a.u. from minimum to maximum (bottom to top). B-C. For visualization purposes spectra are normalized to maxima. Spectra of iBAT (B) and scWAT (C) from chow and HFD-fed mice. Healthy: n = 4, obese: n = 5. D. MSOT image of lipid unmixing from chow and HFD-fed mice. The color bar shows the color coding of MSOT a.u. from minimum to maximum (bottom to top). E-F. Unmixing result of lipid from BAT (E) and scWAT (F). G. MSOT image of TBV unmixing from chow and HFD-fed mice. The color bar shows the color coding of MSOT a.u. from minimum to maximum (bottom to top). H-I. Unmixing result of TBV from iBAT (H) and scWAT (I).

### Combination of MSOT and RSOM visualizes and quantifies accurately exercise-induced morphological changes in vessel density, tissue oxygen saturation, lipid and blood content of dWAT and scWAT

After showing that MSOT accurately detects morphological changes in iBAT and scWAT of DIO mice, we determined whether exercise-induced modulation in adipose tissue mass and function can be monitored by MSOT. As expected HFD fed male and female mice under sedentary conditions exhibited significantly higher body weights compared to sedentary chow diet fed mice. Voluntary wheel running reduced body weights exclusively in HFD-fed male mice to that of chow-fed male mice, indicating a rescue of the DIO phenotype (Figure S4A and B). In addition to iBAT and scWAT, we also analysed dWAT in these cohorts. The absorption spectrum of dWAT was similar to scWAT (Figure 1B, S5). To analyse dWAT in the mouse body, we calculated the percentage of dWAT in the skin, which consists of the epidermis, dermis and hypodermis (dWAT) by MSOT and compared it to the percentage of dWAT in the skin calculated by 2D histology in order to validate our MSOT findings. Using MSOT, we also performed tissue content analysis for TBV to determine blood perfusion in the tissue. In parallel, we performed RSOM imaging to measure dWAT thickness and to visualize the vasculature in the different layers of the skin including dWAT (Fig. 3A). To validate our vascular readouts from MSOT and RSOM, we performed immunohistochemical staining for CD31 on skin tissue slices, a marker commonly used to demonstrate the presence of endothelial cells in histological tissue sections (Bianconi, Herac et al., 2020), and quantified the CD31 positive area (Figure 3A). Our MSOT measurements revealed a significant increase of dWAT percentage in the skin of HFD fed male and female mice compared to chow fed mice (Figure 3A-C; S6A and B). This finding was consistent with results obtained by an independent MRI measurement study (Kasza, Hernando et al., 2016). Voluntary wheel running exercise restored dWAT volume in HFD-fed male mice to normal levels (Figure 3A, B) in contrast to female mice where no significant changes in dWAT volume were detected after exercise (Fig. S6A). Our MSOT findings were consistent with the skin histology of HFD and chow fed mice (Figure 3A, C; Figure S6B). RSOM was able to detect HFD-induced thickening of dWAT as well as the exercise-induced reduction in dWAT thickness in male mice only. (Figure 3A, D). We then measured TBV by optoacoustic methods and validated our findings by analysing the CD31+ area coverage by immunohistochemical staining in the dermis and dWAT (hypodermis) layer of the skin (Figure 3E-G, S7). We found that in dWAT, TBV measured by MSOT was significantly decreased in sedentary HFD-fed mice and this phenotype was rescued by voluntary exercise in male mice only (Figure 3E). This increase in exercise-induced vascular density in HFD-fed male mice was confirmed by our histological staining. Although our histological CD31 based analysis confirmed a reduction in vascular density of HFD-fed mice compared to chow fed mice it did not reach statistical significance as we observed by MSOT (Figure 3F). We assume that this inconsistence is a result of comparing a two-dimensional output (area by CD31+staining) with the three-dimensional output (blood volume by MSOT). RSOM measurements detected a DIO-induced decrease of vessel density and the restoration of the latter by exercise in male mice (Figure 3G). Exercise did not affect vessel density in the dWAT of female mice (Fig S7). In contrast to dWAT (Fig. S6), dermis had no change in CD31+ area coverage measured by histology and vessel density measured by RSOM, indicating an unaltered vascular function in dermis in female DIO mice no matter whether they underwent voluntary running or not (Figure S7). MSOT was not employed for dermis analysis because of the limitation of resolution.

**Figure 3.**
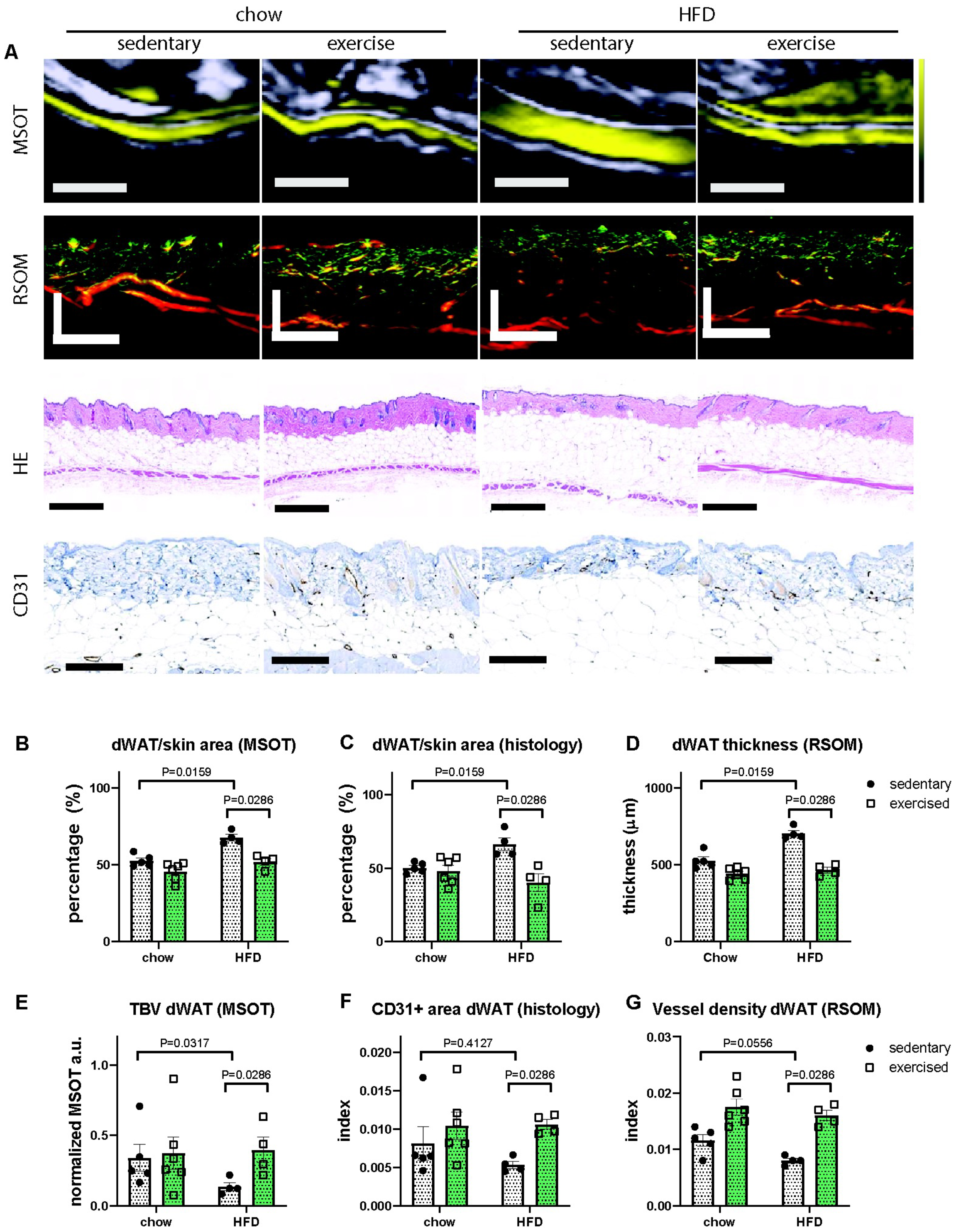
Comparison of dermal adipose tissue (dWAT) characteristic in chow and HFD-fed mice with or without exercise. A. MSOT, RSOM, HE and CD31 staining images from chow and HFD-fed mice with or without exercise. Scale bar: MSOT 2 mm, RSOM 500 μm, HE 500 μm, CD31 200 μm. B-C, Percentage of dWAT in skin calculated from MSOT (B) and histology (C). D. dWAT thickness measured by RSOM. E. TBV in dWAT results measured by MSOT. F. CD31+ area coverage index in whole dWAT. G. TBV in dWAT results measured by RSOM. For data in B, C, E, F, Chow sedentary: n = 5, chow exercised: n = 6, HFD sedentary: n = 4, HFD exercised: n = 4. For data in D and G, each group n = 5.

## Discussion

In this study, we verified that MSOT can visualize and quantify lipid and blood contents in various adipose tissues, including iBAT, scWAT and dWAT of mice. For the first time to our knowledge, we observed *in vivo* that scWAT and iBAT exhibit a decrease in vascularization when DIO occurs. Our non-invasive *in vivo* measurements replicate earlier findings by others using invasive *ex-vivo* and end-point *ex vivo* methods (Graupera & Claret, 2018, Trayhurn, 2013). Our approach allows repetitive and longitudinal non-invasive monitoring of vascular function in adipose tissues undergoing pathophysiological processes *in vivo*, including, but not limited to, DIO associated microvascular disease. In this study, we applied a preclinical version of MSOT modality. Although the clinical MSOT device shares the same principles with the preclinical MSOT device we used herein, the measurement and the quantification procedure need to be adjusted for human subjects, in which the depth of targeted tissue is different from mice.

Here we found that that MSOT can provide a quantitative readout for blood volume in the tissue, but at the same time it does not allow visualization of small vessels due to insufficient resolution. To study the morphological changes in tissue vasculature, RSOM is a better choice of optoacoustic method compared to MSOT. It is important to note that RSOM can only reach about 5 mm depth in mouse tissue. Thus, RSOM is more suitable than MSOT for the study of dWAT, but not of iBAT or scWAT. Besides the visualization of morphological changes in the vasculature of the skin, which cannot be achieved by MSOT, RSOM can also provide quantitative readouts of skin layer thickness, vascular density of different skin layers including dWAT (hypodermis) and dermis. The latter may be of critical importance for the accurate assessment of treatment success in diabetic patients suffering from skin frailty and impaired wound healing. In this study, we only employed one wavelength for RSOM imaging allowing the visualization of blood. Since this wavelength does not allow lipid visualization, the identification of dWAT was based on vessel density, which is much lower in dWAT as compared to the attached dermal layer. However, a multi-spectra method has been recently developed for RSOM technology that will allow visualization of multiple contrasts in the tissue including lipid in future studies (Berezhnoi, Aguirre et al., 2019). By using this multi-wavelength RSOM approach, measurements of dWAT will facilitate a precise skin layer analysis.

Monitoring dWAT vascular function in physiological and pathological conditions, such as hair growth and wound healing, will provide valuable information on morphological changes in these processes, and most importantly provide tools to assess treatment response to vasculature-targeting drugs. For example, there is recent evidence that after depilation, HFD fed mice have a delayed entry into anagen phase in hair growth, in which hair follicles are in contact with dWAT to gain nourishment from dWAT’s blood supply (Zhang et al., 2019). Our observations on dWAT vascular dysfunction under DIO conditions may contribute to further understand as how obesity may affect hair growth.

Clinical applications of optoacoustic imaging are rapidly emerging. Studies using clinical MSOT and RSOM indicated a great potential of these optoacoustic modalities for diagnosing skin diseases, vascular diseases, inflammatory diseases, and cancer (Attia, Balasundaram et al., 2019). However, these applications are mainly taking the advantage of optoacoustic imaging in visualization and quantification of blood content in the tissue and tissue metabolism since the dominant endogenous contrast in most of the tissues is from haemoglobin. In this study, we applied optoacoustic imaging to visualize and quantify lipid and blood content simultaneously to monitor pathological changes in adipose tissues under DIO conditions. Furthermore, we found that voluntary running exercise rescues vascular dysfunction caused by DIO in male mice in contrast to female mice. Our findings set up a base for future clinical studies using optoacoustic-derived vascular readouts from adipose tissues as a biomarker for the personalized monitoring of vascular function in response to stimuli or therapy.

## Methods

### Multi-spectral Optoacoustic Tomography and image analysis

MSOT measurements were performed using a 256-channel real-time imaging system (inVision 256, iThera Medical, Germany). The detailed information of the system has been reported in our previous work(Ntziachristos & Razansky, 2010, Razansky, Buehler et al., 2011). For the measurements in mice, 27 optical wavelengths in the range of 700-960 nm with step of 10 nm were applied to collect multi-spectral optoacoustic signals by using an optical parametric oscillator laser with a 50 Hz repetition rate. The optoacoustic signals were averaged 10 times at each wavelength during data acquisition. For *in vivo* measurements, animals were anaesthetized by continuous inhalation of 2% isoflurane (vaporized in 100% oxygen at 0.8 l/min) and subsequently placed within an animal holder in a supine position relative to the transducer array. The animals were kept into a thin, clear, polyethylene membrane and positioned in the water bath maintained at 34 degrees, which provided acoustic coupling and maintained animal temperature while imaging. The detailed procedure of handing mice in the MSOT imaging system was clearly described in our previous work(Ntziachristos & Razansky, 2010, Razansky et al., 2011). MSOT data were analysed by ViewMSOT software (v3.8, *iThera* Medical, Munich, Germany). MSOT images were reconstructed using the model linear method. For unmixing of Hb, HbO2, Lipid, H2O, and ICG, a linear regression method was used. Each unmixing data point for statistics was averaged from three ROIs in the same subject.

### Histopathology

Adipose tissue specimen were sampled according to established organ sampling and trimming guidelines for rodent animal models (Ruehl-Fehlert, Kittel et al., 2003). The samples were fixed in neutrally-buffered 4% formaldehyde solution for 24 hours and subsequently routinely embedded in paraffin. 3 µm thick sections were stained with haematoxylin and eosin (HE), using a HistoCore SPECTRA ST automated slide stainer (Leica, Germany) with prefabricated staining reagents (HistoCore Spectra H&E Stain System S1, Leica, Germany), according to the manufacturer’s instructions. Histopathological examination was performed by a pathologist in a blinded fashion (*i*.*e*., without knowledge of the treatment-group affiliations of the examined slides).

### RSOM imaging and data analysis

The present study used an in-house portable RSOM imaging system featuring a transducer with central frequency of 50 MHz, which has been described in detail elsewhere (Aguirre, Schwarz et al., 2017). An Onda laser (Bright Solutions, Italy) with dimensions of 19×10×9 cm3 was used to provide light with wavelength of 532 nm. The repetition rate of the laser was 1 kHz, yielding an optical fluence of 3.75 µJ/mm2 under the safety limit. The anaesthetized mouse was placed onto a bed and into a warmed water bath, with the scanned region under the water level and the head above the water level. An optically and acoustically transparent plastic membrane was affixed using surgical tape on the mouse skin at the scanned region. The scanning head containing the laser output and transducer was brought close to the membrane to position the focal point of the ultrasound detector slightly above the skin surface and thereby maximize detection sensitivity. The scanning head contained water as coupling medium. Two mechanical stages (PI, Germany) were used to scan the RSOM head. The laser and controller of the mechanical stages were both stored inside a plastic case, which ensured laser safety. The scanning field of view is 4×2 mm^2^ with step size 7.5 µm in the fast axis and 15 µm in the slow axis. The total scanning time of one measurement took about 70 s. For image reconstruction, optoacoustic signals were separated into lower (10–40 MHz, red) and higher (40–120 MHz, green) frequencies to distinguish larger (diameter of 50 to more than 100 µm) and smaller (diameter of 10–40 µm) vessels, respectively. This bandwidth separation was performed for all RSOM dataset using the same method by using the same frequency ranges, meaning that larger (red encoded) and smaller (green encoded) vessels represent the same size range throughout all mice measurements. The two reconstructed images *R*_*low*_and *R*_*high*_ corresponded to the low- and high frequency bands. A composite image was constructed by fusing *R*_*low*_into the red channel and *R*_*high*_ into the green channel of a same RGB image. The detail process has been introduced in our previous work (Aguirre et al., 2017).

To compute dWAT thickness, RSOM images were first flatted based on our surface detection approach (Schwarz, Garzorz-Stark et al., 2017). The reconstructed volume of selected frequency band (10-40 MHz) was split into four stacks with 0.5 mm thickness along the slow scanning axis. Then, the dermis layer in the MIP image of each stack were automatically segmented by graph theory and dynamic programming-based approach (Chiu, Li et al., 2010). The thickness of the dermis layer was calculated as the average width of the four segmented boundaries. The dermis layer was segmented as starting from the bottom boundary of the dermis layer and further extending 1 mm depth. In the 4×2 mm scanning region, the vessel density in the segmented dermis layer was calculated as *N* × *dV*, where N represents the number of voxels with intensity above 20% of the maximum voxel intensity, and *dV* is the voxel volume.

### Animal studies

Age-matched mice (Jackson laboratory; strain # B6(Cg)-*Tyr*^*c-2J*^/J) were kept at 22 ± 2 °C with constant humidity (45–65%) and a 12-h light–dark cycle and with ad libitum access to food and water. After a seven-day acclimation phase, mice were randomly divided into chow (Altromin 1310; 14 kcal% fat, 59 kcal% carbohydrates) and high fat-diet (HFD; D12331; 58 kcal% fat and 25.5 kcal% sucrose, Research Diet, New Brunswick, NJ, USA) fed groups. Each chow and HFD groups were further randomly divided into either sedentary control group or exercise group. The exercise group had free access to voluntary wheels within their home cage. The wheel running profile (Figure S8) which includes time, duration, and speed was monitored using a commercially available wifi wheel running system (Low-Profile Wireless Running Wheel, Med associates inc, St. Albans, VT 05478, US). To avoid social stress, two mice per cage were housed throughout the duration of the study. At the end of the study, mice were euthanized with an overdose of ketamine and xylazine and blood and organs were collected. The animal studies were approved and conducted in accordance with the Animal Ethics Committee of the government of Upper Bavaria, Germany.

### Statistics

Data were analyzed using GraphPad Prism (v. 8.4.2; GraphPad Software, La Jolla, CA). All data presented as mean ± SEM unless otherwise stated. Group size (n) is indicated for each experiment in figure legends. Sample size was determined by the statistical report for the animal experiment application in accordance with the principle for replacement, refinement or reduction of the use of animals in research and approved by the Animal Ethics Committee of the government of Upper Bavaria, Germany. Student’s t-test was used for comparisons of two independent groups. One-way ANOVA followed by Tukey’s post hoc test was used for comparing more than two independent groups. A p-value of < 0.05 was considered as statistically significant. Significant digit: *P< 0.05, **P< 0.01, *** P< 0.001.

## Acknowledgements

The authors would like to thank Uwe Klemm for technical assistance, Doris Bengel for support in animal experiments. We are grateful for the support from our husbandry staff at the Research Unit Comparative Medicine and the staff of the Core Facility Pathology and Tissue Analytics of Helmholtz Munich. This work was supported by the German Research Council (DFG) grants HO 2286/3-1 to S.M.H. and NT 3/32-1 to V.N. as part of the Research Unit FOR 5298 (iMAGO) as well as the DFG Gottfried Wilhelm Leibniz Prize 2013 (NT 3/10-1), the European Research Council (ERC) under the European Union’s Horizon 2020 research and innovation programme under Grant Agreement No. 694968 (PREMSOT) and the Helmholtz-Gemeinschaft Deutscher Forschungszentren (HGF)/ExNet project “Innovative Intelligent Imaging” (i3-Helmholtz) to V.N.

## Author contributions

SH conceived the study, designed, and performed the experiments, analysed the data and wrote the manuscript. HH conceived the study, designed, and performed the experiments, analysed the data, and wrote the manuscript. RZT performed the experiments, analysed the data, and wrote the manuscript. SG performed the in vivo experiments. PA performed the in vivo experiments. ACS conceived the study. SMH conceived the study and wrote the manuscript. VN conceived the study and wrote the manuscript.

## Disclosure and competing interest statement

V.N. is an equity owner and consultant for iThera Medical GmbH, Munich, Germany

## The paper explained

### Problem

Microcirculatory dysfunction has been observed in dermal and subcutaneous fat layers of obese humans and has been proposed as an early prediction marker for cardio-metabolic disease progression. In-vivo visualization and longitudinal monitoring of microvascular remodeling in these tissues remains challenging.

### Results

Using a combination of non-invasive optoacoustic imaging methods MSOT and RSOM provided accurate and longitudinal non-invasive monitoring of microcirculatory dysfunction in dermal and subcutaneous adipose tissues *in vivo* in a mouse model for diet-induced obesity (DIO). Non-invasive optoacoustic imaging methods MSOT and RSOM visualized accurately revascularization in dermal and subcutaneous fat after voluntary wheel running in DIO mice.

### Impact

Our findings indicate that optoacoustic-derived vascular readouts from dermal and subcutaneous adipose tissues may be used as a personalized biomarker for monitoring microcirculatory function in cardio-metabolic disease.

## Data Availability

The primary MSOT and RSOM data that support the findings of this study are not openly available due to reasons of size (1.3TB) and are available from the corresponding author upon reasonable request.

## Abbreviations

dWAT: Dermal white adipose tissue
scWAT: Subcutaneous white adipose tissue
iBAT.: Interscapular brown adipose tissue
MSOT: Multi-spectral optoacoustic tomography
RSOM: Raster-scanning optoacoustic mesoscopy
DIO: Diet-induced obesity
MRI: Magnetic Resonance Imaging
CT: Computed Tomography
BAT: Brown adipose tissue
WAT: White adipose tissue
HFD: High fat diet
Chow Standard diet
Hb: deoxy-haemoglobin
HbO_2_: oxy-haemoglobin

## Figure legends

**Figure S1.**
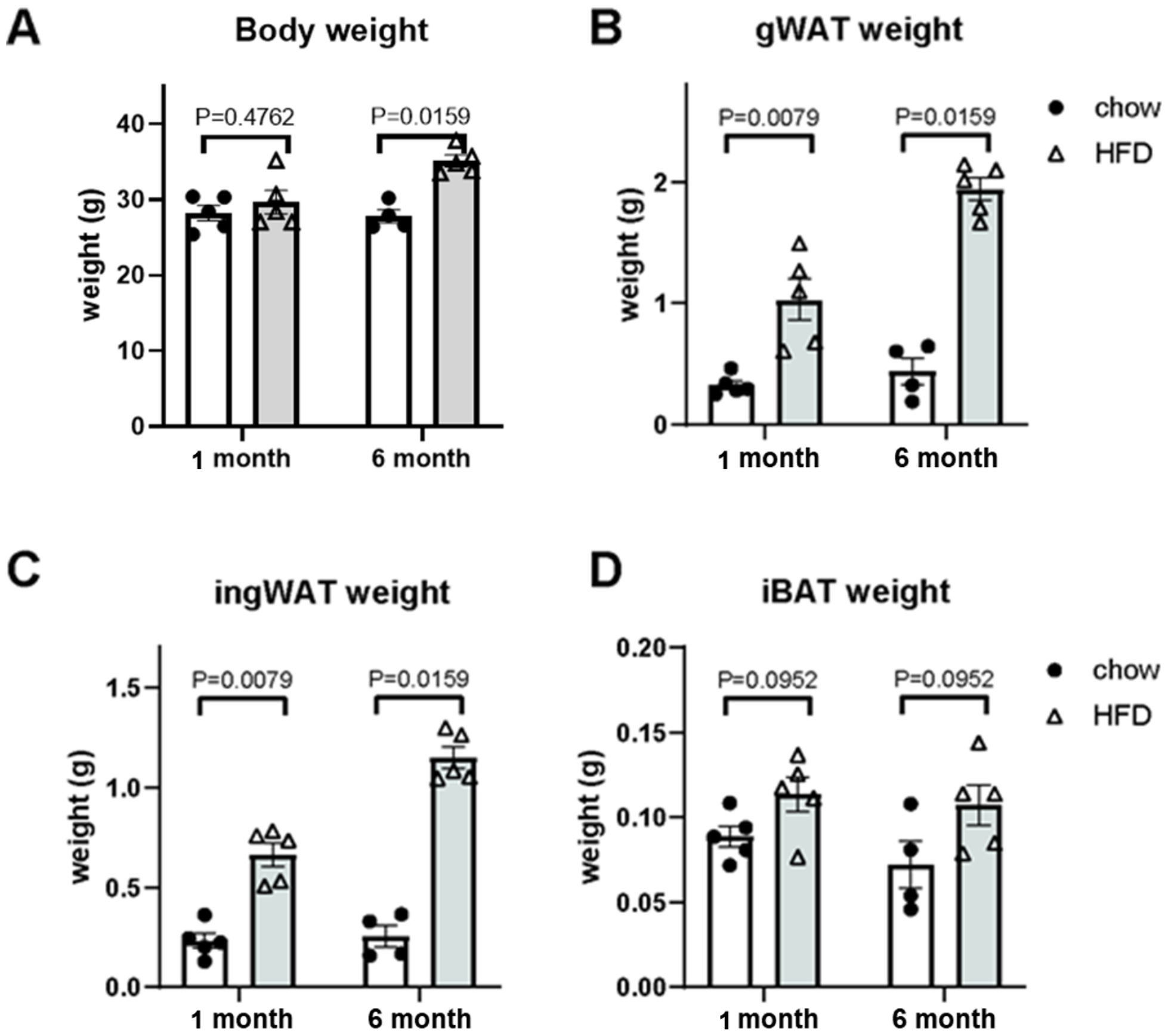
Body weight and the weights of gonadal white adipose tissue (gWAT), inguinal white adipose tissue (ingWAT), interscapular brown adipose tissue (iBAT) from male mice fed with chow and high fat diet (HFD). A. Body weight of mice after 1 month or 6 months feeding with chow and HFD. B-D. The weights of gWAT (B), ingWAT (C), iBAT (D) of mice after 1 month or 6 months feeding with chow or HFD. For data in all panels: chow n = 4 - 5.

**Figure S2.**
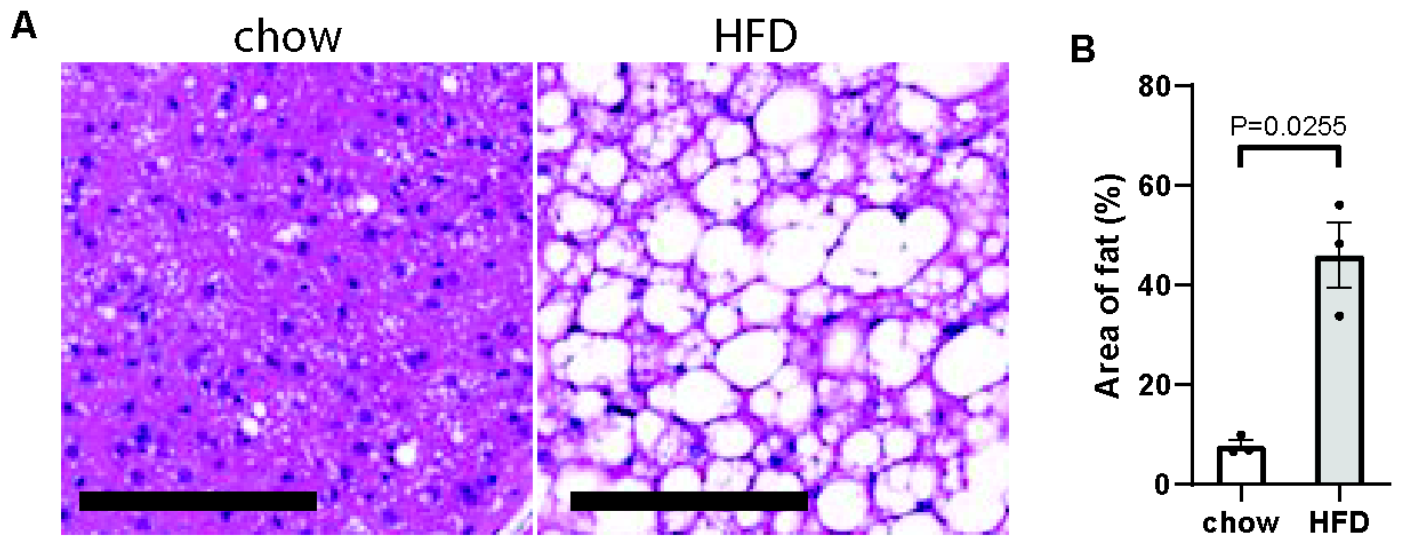
Whitening of brown adipose tissue in high fat diet (HFD) fed mice. A. HE staining of brown adipose tissue from mice fed with chow and HFD. Scale bar: 100 μm. B. Quantification of fat area coverage in HE staining images. For each group: n = 3.

**Figure S3.**
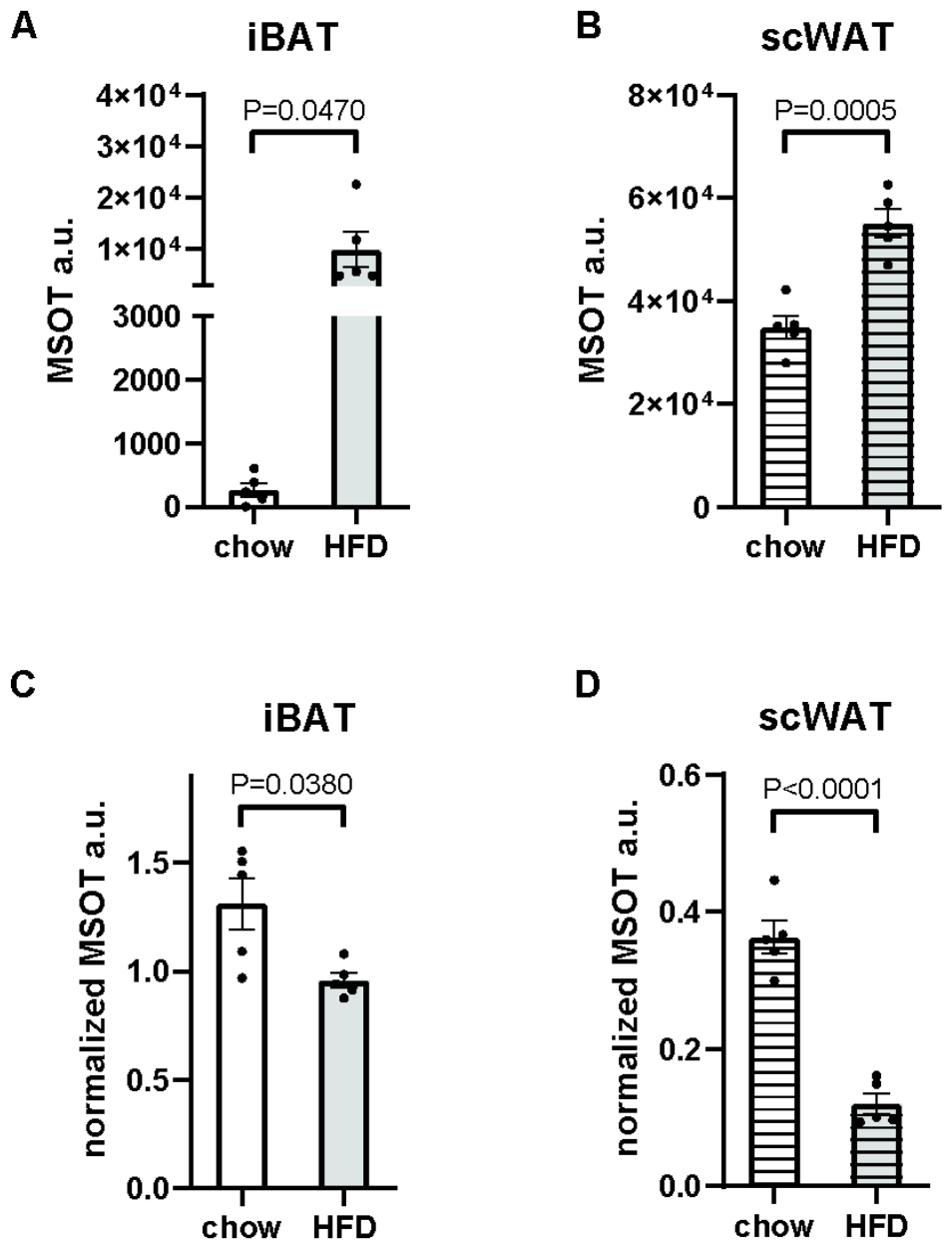
MSOT unmixing results of lipid and total blood volume (TBV) of interscapular brown adipose tissue (iBAT) and subcutaneous white adipose tissue (scWAT) in male mice after 1 month of HFD feeding. A-B. Lipid unmixing results from iBAT (A) and scWAT (B). C-D. TBV unmixing results from iBAT (C) and scWAT (D). For all groups in each panel: n = 5.

**Figure S4.**
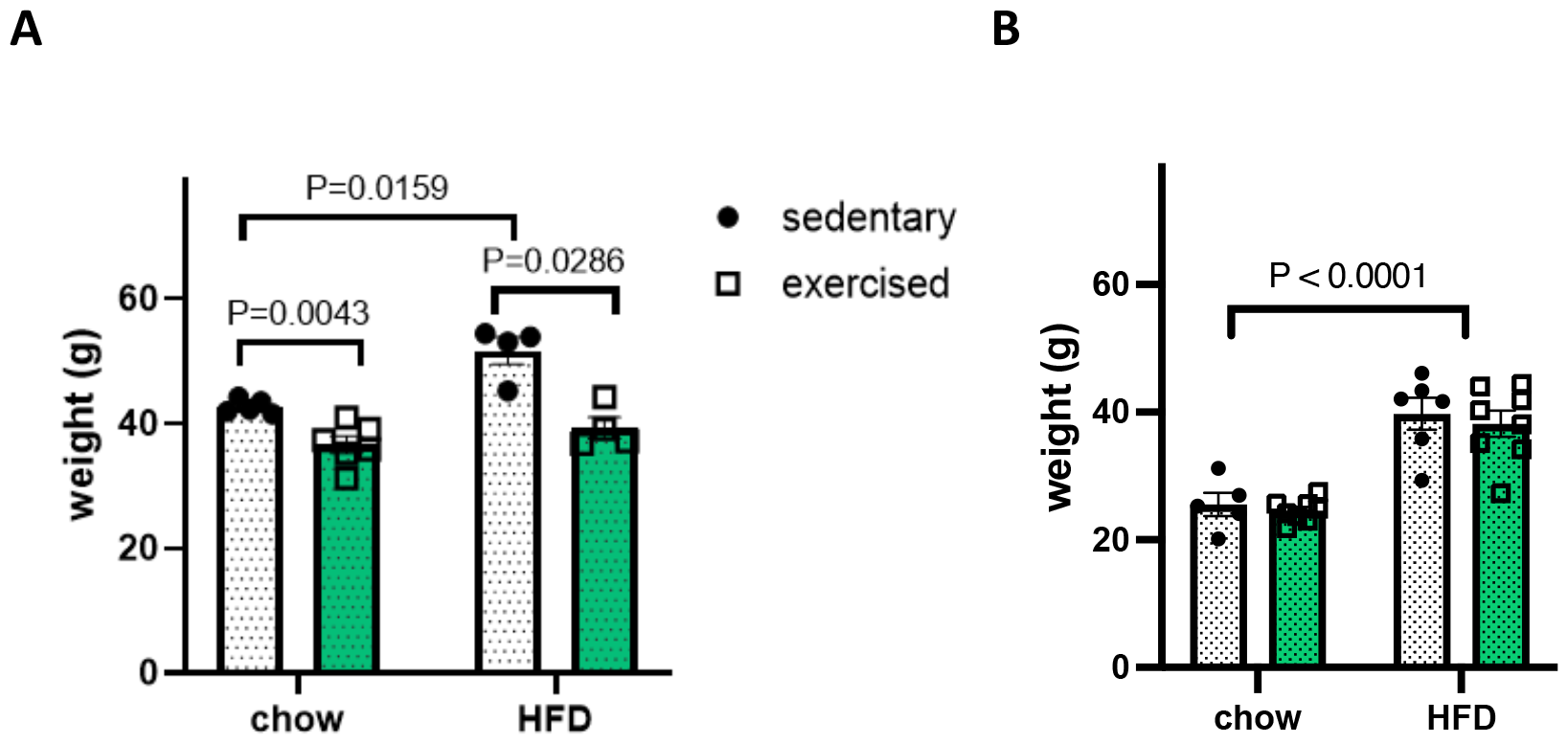
Body weights of male mice (A) and female mice (B) fed with chow or high fat diet (HFD) with or without voluntary running wheel exercise. N = 4-8.

**Figure S5.**
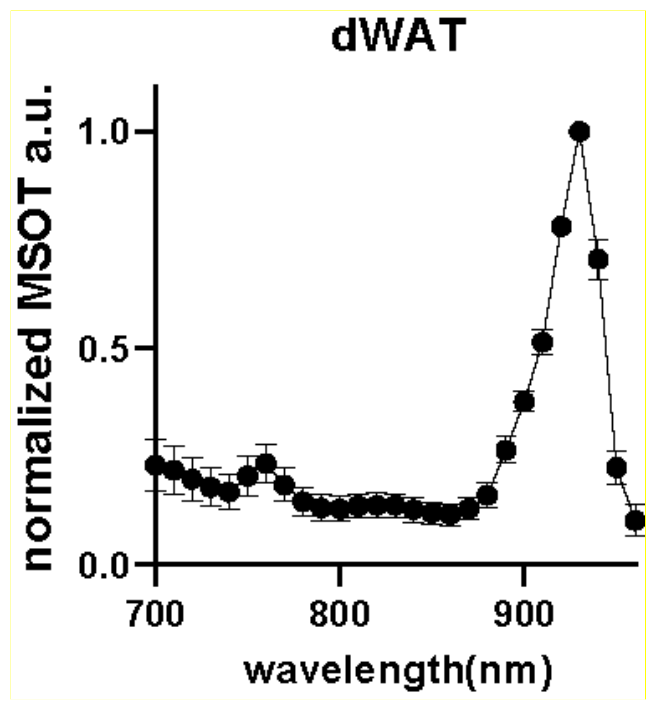
Normalized spectra of dermal white adipose tissue (dWAT) from normal mice. n = 5.

**Figure S6.**
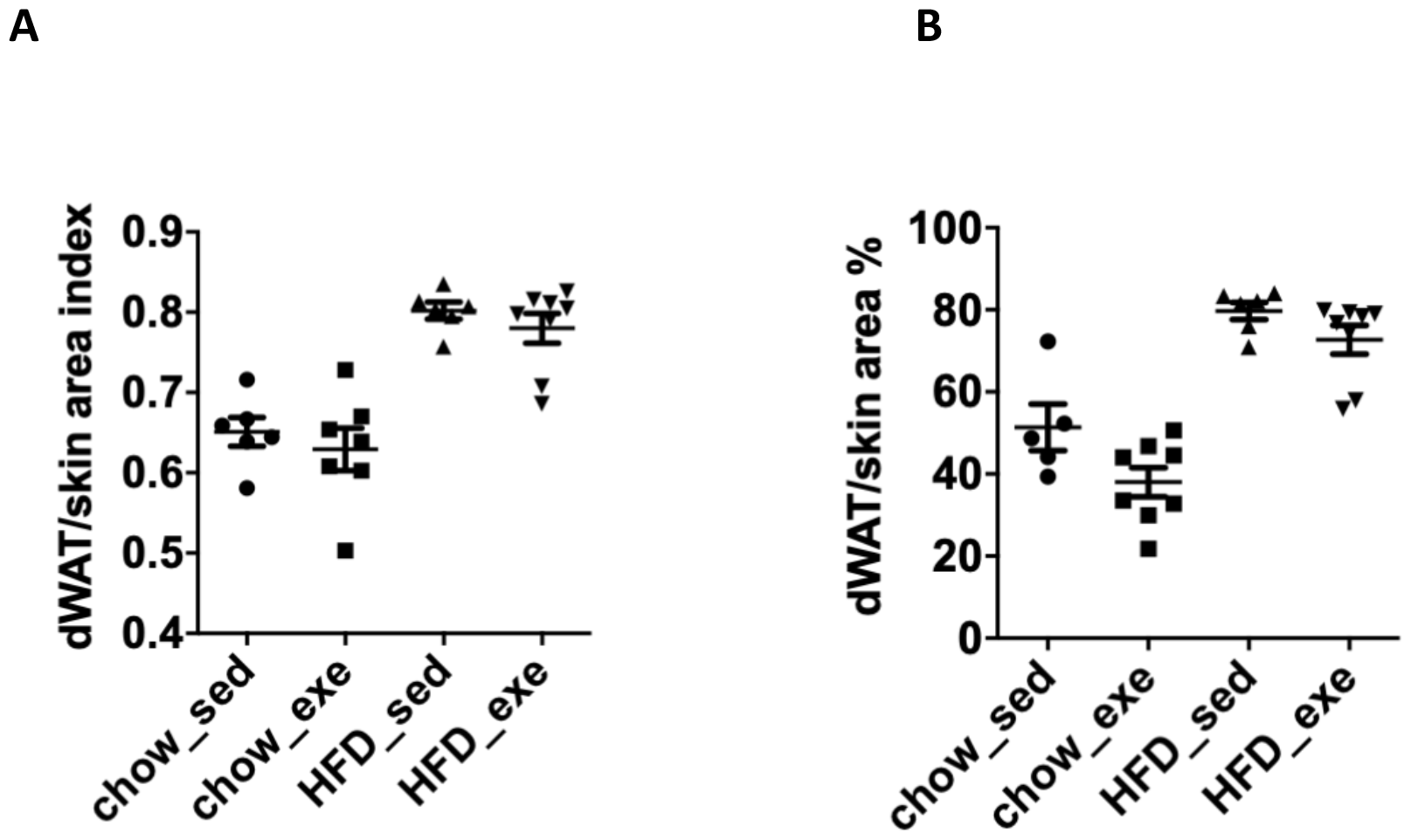
Comparison of dermal adipose tissue (dWAT) characteristic in chow and HFD-fed female mice with or without exercise: Percentage of dWAT in skin calculated from MSOT (A) and histology (B).

**Figure S7.**
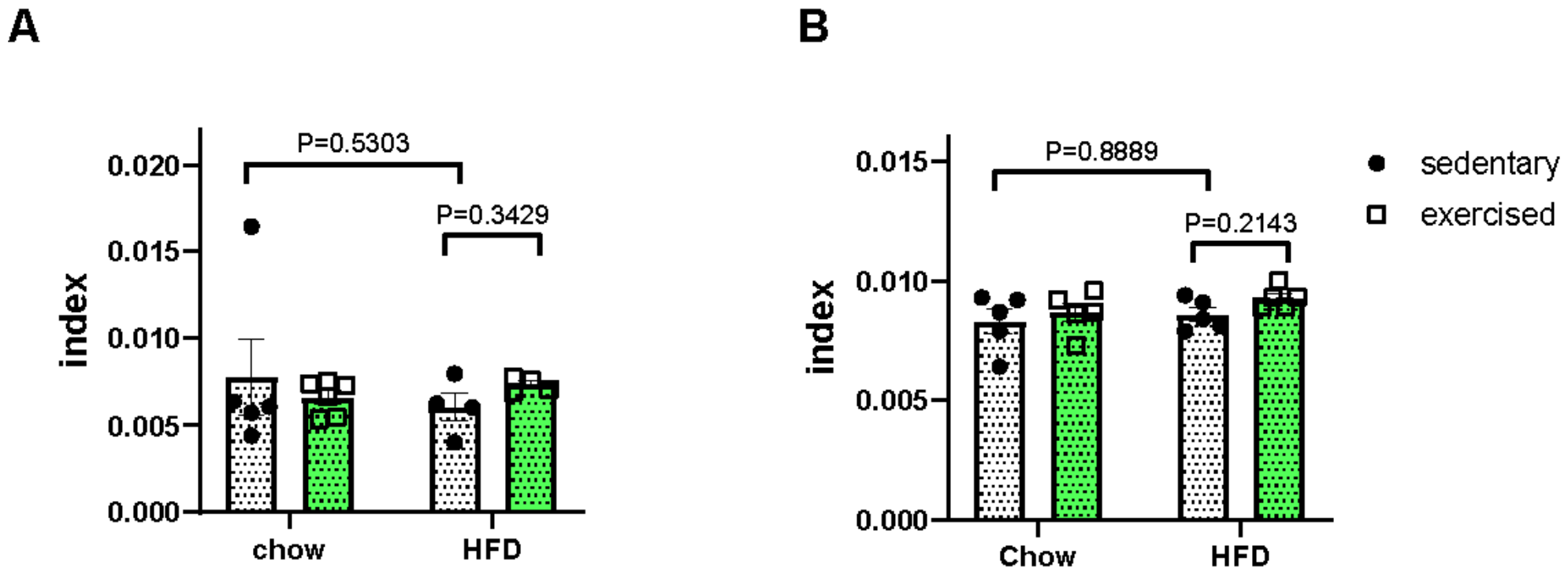
Vessel density in dermis of female mice fed with chow or high fat diet (HFD) with or without exercise. CD31+ area coverage in dermis of mice fed with chow or high fat diet (HFD) with or without exercise (A). RSOM result of total blood volume (TBV) in dermis of mice fed with chow or high fat diet (HFD) with or without exercise (B). For each group: n = 5

**Figure S8.**
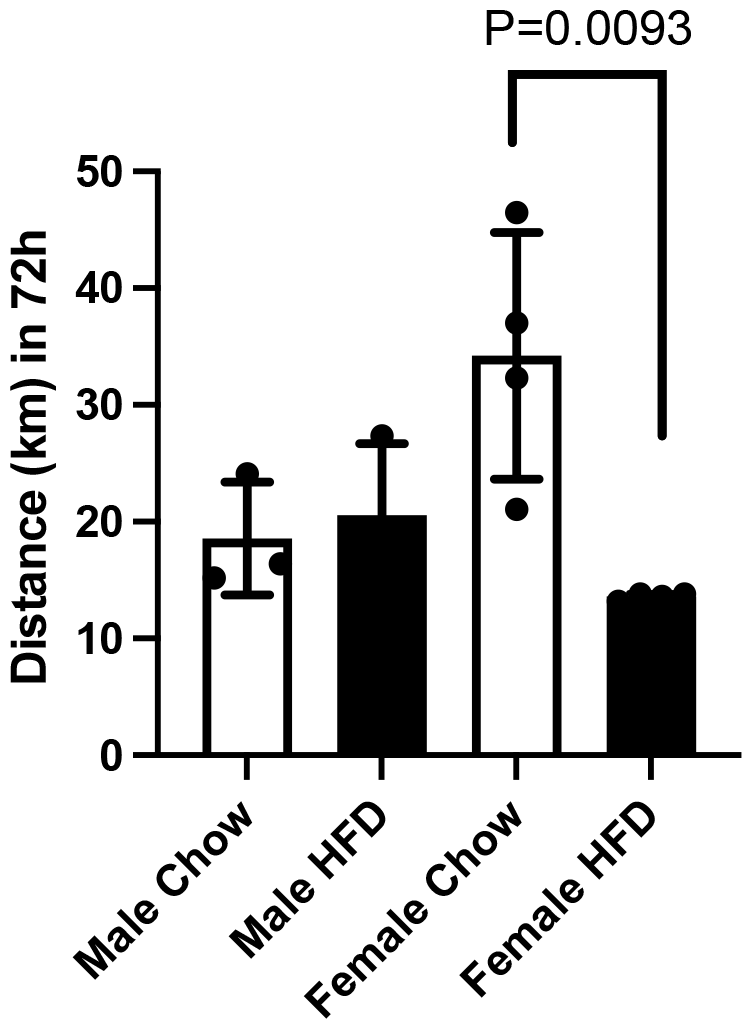
Running distance of male and female mice fed chow and HFD during over 72 hours (n = 6 - 8; 2 mice per cage).

